# Aperiodic slope reflects glutamatergic tone in the human brain

**DOI:** 10.1101/2025.05.15.654118

**Authors:** Aislin A. Sheldon, Hannah R. Moser, Kamar S. Abdullahi, Karly D. Allison, Carter B. Mulder, Samantha A. Montoya, Scott R. Sponheim, Małgorzata Marjańska, Michael-Paul Schallmo

## Abstract

Excitatory and inhibitory neural processes are essential for every aspect of brain function, but current non-invasive neuroimaging methods to study these in the human brain are limited. Recent studies which separate oscillatory and aperiodic components of electrophysiological power spectra have highlighted a relationship between aperiodic activity and functional brain states. Studies in both animal models and humans suggest that the aperiodic slope of electrophysiological power spectra reflects the local balance of excitatory:inhibitory (E:I) synaptic transmission. Aperiodic slope varies across individuals, brain states, and clinical populations, which may reflect important differences in E:I balance. However, there is currently a lack of evidence linking aperiodic slope to other measures of excitation and inhibition in the human brain. Here, we show that flatter (less steep) aperiodic slopes from human electroencephalography (EEG) are associated with higher concentrations of the excitatory neurotransmitter glutamate measured with 7 tesla magnetic resonance spectroscopy (MRS) in the occipital lobe at rest. This suggests that individual differences in aperiodic neural activity reflect cortical glutamate concentrations, providing important insight for understanding changes in neural excitation across brain states and neuropsychiatric populations (e.g., schizophrenia) where glutamatergic function may differ. Our results support the use of aperiodic slope as a non-invasive marker for excitatory tone in the human brain.

## Introduction

Neural processing in the brain requires a balance of excitation and inhibition (E:I) to facilitate neurotransmission within and across brain regions. As the brain is a highly recurrent, densely interconnected system, this balance is needed to prevent over-excitation, which may cause excessive neural activity resulting in seizures, or over-inhibition, which may prevent the necessary propagation of information via action potentials. E:I balance is thought to differ across brain states, such as during sleep versus wakefulness ^1,2^, and between processing of different sensory stimulus conditions ^3^, as well as varying across individuals and clinical populations ^4,5^. The excitatory and inhibitory properties of the nervous system can be interrogated directly in animal models using invasive tools such as optogenetics and patch clamp electrophysiology ^3,6,7^. However, studies of the human brain primarily rely on non-invasive neuroimaging techniques to measure neural activity. Therefore, there is a need to develop tools to investigate excitation and inhibition non-invasively in the human brain.

One technique for examining E:I balance in the human brain, which has gained interest in recent years, uses analytic tools to separate periodic (i.e., oscillatory) activity from aperiodic (i.e., non-oscillatory) components of the power spectrum measured during electroencephalography (EEG) ^8^. EEG measures changes in voltage driven primarily by postsynaptic potentials of pyramidal cells, the major input-output neurons in the cortex. The aperiodic component of the EEG power spectrum exhibits a 1/f-like pattern, whereby an inverse relationship exists between frequency and power, with more power present at lower frequencies and less power at higher frequencies ^8,9^. The slope of this aperiodic component was originally thought to reflect neural noise, but recent work in animal models has suggested that the aperiodic slope reflects the local balance of E:I synaptic transmission ^10^, with less steep slopes (i.e., flatter, with similar power across frequencies) observed under conditions of relatively higher excitation, and steeper slopes with relatively higher inhibition. One explanation for this may be the comparatively faster dynamics of excitatory postsynaptic currents driven by glutamatergic AMPA receptors, relative to slower inhibitory postsynaptic currents from GABA_A_ receptors ^10^. Consistent with this, studies in humans have shown changes in the aperiodic slope across task conditions ^11,12^, age ^13–16^, and in different clinical populations ^17,18^ in which E:I functions are thought to differ.

Another neuroimaging tool that provides insight into excitation and inhibition in the human brain is magnetic resonance spectroscopy (MRS) which can be used to non-invasively measure the concentrations of neural metabolites such as glutamate and γ-aminobutyric acid (GABA), which are the primary excitatory and inhibitory neurotransmitters in the human brain, respectively. MRS is thought to measure glutamate present in the cytosol of neurons (including pyramidal cells) and glial cells ^19^. Glutamate and GABA levels from ‘static’ or ‘resting state’ MRS are likely more reflective of oxidative energy metabolism ^19^, or excitatory and inhibitory tone ^20,21^, rather than levels of neurotransmitters directly involved in synaptic transmission. Therefore, individuals or clinical populations with higher baseline levels of glutamate or GABA in a given region may have a greater potential for cortical excitation or inhibition, respectively ^22^. Numerous MRS studies have shown that resting levels of glutamate and / or GABA may differ across brain regions, individuals, and clinical populations ^23–25^. Further, studies using functional MRS have found that concentrations of glutamate and GABA change in response to sensory input ^26–32^, and changes to cognitive demands ^33,34^; see Mullins ^35^ for a meta-analysis and Koolschijn et al. ^36^ for a review.

The current study consisted of two separate experimental sessions: 1) EEG measurements of the aperiodic slope of the power spectrum during two rest conditions (eyes closed and eyes open), and 2) MRS measurements of glutamate and GABA within the occipital cortex during eyes open fixation. In order to investigate whether individual differences in aperiodic slope are related to metabolites involved in excitation and inhibition, we tested whether aperiodic slope correlated with glutamate and GABA concentrations from MRS. Based on the hypothesis that aperiodic slope is indicative of neural E:I balance, we predicted that higher (more steep) aperiodic slopes would exhibit a negative correlation with concentrations of the excitatory neurotransmitter glutamate and a positive correlation with concentrations of the inhibitory neurotransmitter GABA in occipital brain regions. Consistent with our hypothesis, we observed that individuals with a higher glutamate concentration had a less steep aperiodic slope. However, GABA concentrations were not significantly correlated with aperiodic slope. Overall, our findings indicate that individual differences in aperiodic EEG slope may be driven, in part, by differences in glutamate levels, and this could provide a useful non-invasive marker of excitatory tone in the human brain.

## Methods

### Participants

We collected EEG data from 35 adult participants without a history of significant neurological conditions (see Supplemental Methods for full exclusion criteria); of these, 27 participants also completed the MRS protocol. EEG and MRS data were acquired in separate sessions on different days, with MRS typically acquired second (mean number of days = 56, *SD* = 57, median = 25). Of the individuals who had both EEG and MRS data, 1 participant was excluded for data quality reasons (see data processing and analysis section), therefore our final sample for the combined analyses was 26 participants (demographics are shown in Supplemental Table 1). All participants provided written informed consent and were compensated $20 per hour. Ethical approval was granted by the Institutional Review Board at the University of Minnesota and all procedures complied with the Declaration of Helsinki.

### Apparatus

#### EEG

EEG data were recorded from 32 Ag/AgCl electrodes (BrainVision ActiCap) in a custom 10-10 system ^37–39^ montage with a higher density of electrodes over the posterior scalp, as pictured in Figure 1C. Additional peripheral bipolar electrodes were applied to the face and used to record horizontal and vertical eye movements (see Supplemental Methods for further details).

**Figure 1.**
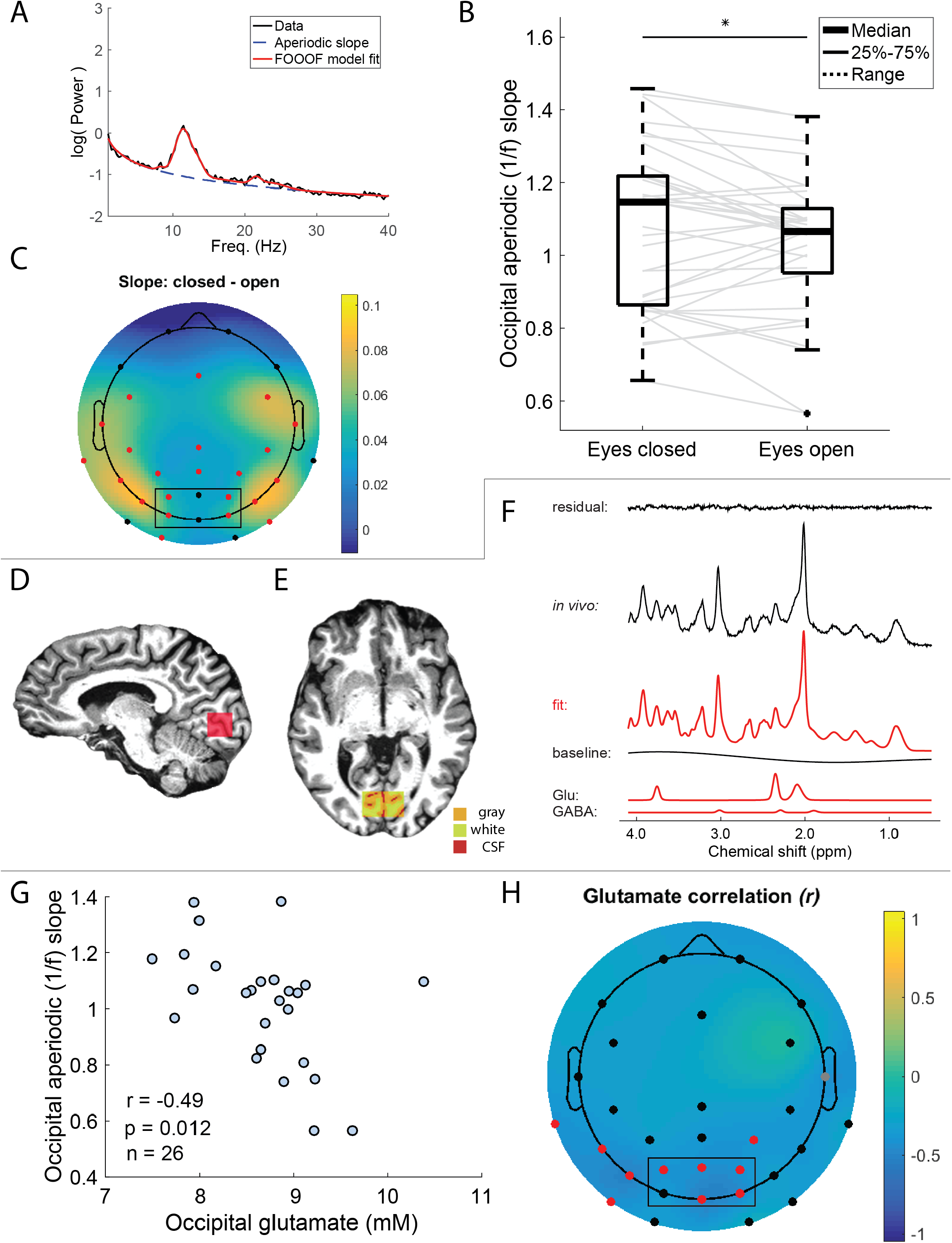
A) FOOOF model applied to eyes open resting data from an example participant. B) Difference between the aperiodic 1/f slope for eyes closed versus eyes open from the 6 occipital electrodes (O1, Oz, O2, PO3, POz, PO4; black box). C) Topoplot shows the difference in aperiodic slope between eyes open and eyes closed conditions. Red dots represent electrodes with a significantly greater slope with eyes closed (cluster corrected *p* < 0.05, one-tailed *t*-test). D) MRS VOI position: sagittal view with VOI in red. E) Segmentation of axial MRS VOI content into GM (orange), WM (yellow), and CSF (red). F) Example MR spectrum from an individual participant (black) plus LCModel fit (red), residual and baseline. The bottom rows show glutamate and GABA signals, as fitted by LCModel. G) Correlation between eyes open aperiodic 1/f slope from 6 occipital electrodes (O1, Oz, O2, PO3, POz, PO4; black box) and glutamate concentration. H) Topoplot showing the correlation between occipital glutamate concentration and eyes open aperiodic slope measured at each electrode. The interpolated color plot over the scalp shows the *r* value. Points show the electrode locations, red indicates a significant correlation (cluster corrected *p* < 0.05), gray indicates electrodes that did not pass cluster correction (uncorrected *p* < 0.05).

#### MRS

Proton magnetic resonance spectroscopy (^1^H MRS) data were acquired at 7 T with a Siemens MAGNETOM scanner. We used a custom-built radio frequency head coil with one surface ^1^H quadrature transceiver for the occipital cortex (Figure 1D). We acquired *T*_1_-weighted and *T*_2_-weighted anatomical images at 3 T on a whole-body Siemens PRISMA scanner with a 32-channel receive-only head coil (see Supplemental Methods for further details).

#### Experimental protocol EEG

EEG data were collected in a darkened room during a 4-minute resting paradigm in E-Prime (v3, Psychology Software Tools). The task consisted of two conditions: eyes open or eyes closed (two 1-minute blocks of each in the order: closed, open, closed, open), with a recorded audio cue (“close your eyes” or “open your eyes”) delivered via earphones at the start of each block (see Supplemental Methods for further details).

#### MRS

The following methods conform to the consensus for MRS reporting standards ^40^. In a separate study visit from the EEG, we acquired MRS data at 7 T with a stimulated echo acquisition mode (STEAM) sequence with echo time (TE) = 8 ms (see Supplemental Methods for further details). Data were acquired with 3D outer volume suppression that was interleaved with variable power and optimized relation delay (VAPOR) water suppression ^41^. We used a visual stimulus (see below for parameters) to functionally localize visual responses in the occipital lobe. We placed the MRS volume of interest (VOI; 30 x 30 x 18 mm^3^) over the mid-calcarine to maximally cover the occipital activation while avoiding the sagittal sinus and dura. Average VOI placement across participants is shown in Supplemental Figure 1. Gradient echo functional magnetic resonance imaging (fMRI) data were acquired for this localizer scan (see Supplemental Methods for further details). We ensured adequate water suppression prior to MRS data collection. We acquired 60 FIDs at rest which were collected as part of a larger scanning run which included an additional 60 FIDs during presentation of visual stimuli; each participant completed 2 to 4 runs (120 to 240 FIDs at rest). We acquired a water reference scan (1 FID, transmitter frequency = 4.7 ppm) for eddy current correction and to quantify metabolite concentrations relative to water. FAST(EST)MAP ^42,43^ was used for B_0_ shimming to obtain a water linewidth of ≤ 15 Hz; data were not collected if the linewidth was larger than this value.

We acquired *T*_1_-weighted and *T*_2_-weighted anatomical images for MRS VOI registration and to correct for tissue fractions. *T*_1_-weighted anatomical images were also acquired at 7 T during the MRS experiment and later aligned to the whole-brain 3 T *T*_1_ images.

#### MRS task

MRS data were collected while participants performed a parafoveal fixation task designed to maintain vigilance and control visual attention. The same task was presented during the water reference scan. Both ‘rest’ (blank background) and ‘stimulus’ (contrast reversing grating) runs were collected; only rest runs were used in the current analyses (see Supplemental Methods for further details). For inclusion in the current analysis, participants needed to have completed at least two (and up to four) 5-minute MRS rest runs. The same stimuli were used for the functional localizer fMRI scan, which facilitated MRS VOI placement as described above.

#### Data processing and analysis EEG

EEG data were preprocessed using EEGLab (v2020.0; ^44^) in MATLAB (2021b). Continuous data were down-sampled to 250 Hz, then high-pass filtered at 0.05 Hz and notch filtered (58 - 62 Hz) to remove AC electrical noise, using a Hamming windowed sinc finite impulse response filter. Electrodes with an impedance value > 10 kΩ during data collection were excluded (average 0.43 channels per participant) prior to artifact correction via independent component analysis. Individual components were visually inspected and components containing predominantly ocular, muscular, or cardiac artifacts were removed. Excluded channels were then replaced via spherical interpolation. Data were re-referenced to the mean of all scalp electrodes. Power spectral density (PSD) was calculated as follows using Welch’s method (*pwelch* function in MATLAB; ^45^). Data from each 60 s task condition block were epoched into 4 s windows with 50% overlap and 256 discrete Fourier transform points per window, with the PSD from each window averaged across the entire 60 s block. First, we calculated PSD from EEG data averaged across a group of 6 occipital channels (O1, Oz, O2, PO3, POz, PO4; black box in Figure 1C) that are expected to reflect neural signals from the medial occipital lobe (i.e., the same general location as the MRS voxel; Figure 1D). These electrodes were chosen *a priori*.

Second, we calculated the PSD for each of the 32 electrodes separately. In both cases, we calculated the mean PSD for each condition (eyes closed and eyes open) across the two runs. We quantified the slope of the aperiodic (1/f-like) component of the EEG signal for each average PSD within the 1 - 40 Hz range using the default parameters for detecting oscillations with the “fitting oscillations and one over *f*” (FOOOF) toolbox ^8^ in Python (Anaconda 4.9.2; Figure 1B), with the equation:

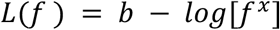

where *f* is the frequency, *x* is the aperiodic exponent (slope on a logarithmic axis), and *b* is the offset (intercept). The goodness-of-fit (*R*^2^ value) for all datasets was above 0.95.

#### MRS

MRS data were preprocessed in MATLAB (2014a) using the *matspec* toolbox (github.com/romainVala/matspec) which included eddy current correction, frequency and phase correction (see Supplemental Methods for further details).

Metabolite concentrations were quantified using the linear combination model (LCModel) version 6.3-1N ^46^. An example spectrum is shown in Figure 1F (VOI shown in red in Figure 1D). We used a basis set which included 18 metabolites (see Supplemental Methods for further details). The macromolecule signals included in the basis set were measured using inversion-recovery experiments ^47^ on the same scanner performed in the occipital cortex (same VOI) of 4 adult participants ^48^. A stiff baseline (spline constraint parameter DKNTMN = 5), and no soft constraints on metabolite concentrations were used to fit spectra ^49–51^. Data were averaged for each individual rest run before fitting (60 FIDs). Metabolite concentrations were scaled relative to water, taking into consideration the *T*_1_ and *T*_2_ of water, and these values are reported in Supplemental Table 2.

FreeSurfer (version 5.3.0) ^52^ was used to generate individual surface model tissue segmentations (Figure 1E) for gray matter (GM; orange in Figure 1E), white matter (WM; yellow in Figure 1E) and cerebrospinal fluid (CSF; red in Figure 1E) from the *T*_1_- and *T*_2_-weighted anatomical images acquired at 3 T.

We used MATLAB (2016a) to define multiple post-hoc MRS data quality metrics based on visual data inspection. Specifically, we plotted histograms to detect outliers and used these to set threshold values for spectral linewidth (≤ 6) and signal-to-noise ratio (SNR; ≥ 30) from LCModel for each individual MRS run. We additionally set a threshold for the minimum response rate across trials for the fixation task during MRS (≥ 85%), and removed any runs for which values were outside these thresholds. Metabolite values for the remaining runs were then averaged. Participants’ data were removed if they had less than two runs retained. Omitting these data quality exclusions did not qualitatively affect our pattern of results (data not shown).

#### Statistical analysis

Statistical analyses were performed in MATLAB (2016a). We used paired *t*-tests to compare slopes between the eyes open and eyes closed conditions. A one-tailed test was chosen to examine our *a priori* hypothesis that slope is less steep with eyes open, based on previous findings ^11^.

We used Spearman rank correlations to quantify relationships between aperiodic EEG slopes across 6 occipital electrodes, and the concentration of glutamate, GABA, and NAA in the occipital lobe from MRS. We used a Bonferroni correction to adjust *p*-values for these 3 multiple comparisons. We also repeated these 3 correlations using slope values from each EEG electrode separately. In this case, we used a cluster correction method to correct for multiple comparisons (see Supplemental Methods for further details). We visualized the correlation *r*-values across electrodes using the *topoplot* function in EEGLab.

To examine the potential role of data quality in our results, we also quantified Spearman correlations between aperiodic slope and MRS SNR, spectrum linewidth, and the area of the unsuppressed water peak (each from LCModel). We also examined whether demographics or other factors might account for the observed results (see Supplemental Methods for further details).

## Results

We measured EEG power spectra during rest in 35 adult participants. We used the FOOOF toolbox to quantify the aperiodic slope of the power spectra, and compared slopes for eyes closed vs. eyes open rest conditions. As expected, we found steeper aperiodic slopes with eyes closed vs. open, both for data averaged across 6 posterior electrodes (O1, Oz, O2, PO3, POz, PO4; *t*_34_ = 2.2, *p* = 0.032), and for the majority of individual electrodes (22/31; cluster corrected *p* = 0.005; Figure 1A and C). Less steep slopes with eyes open is consistent with higher excitation and increased E:I ^11^.

We examined the association between aperiodic slopes (with eyes open) and concentrations of the excitatory neurotransmitter glutamate measured in 26 participants in the visual cortex during a fixation task. Less steep aperiodic slopes from occipital electrodes (box in Figure 1H) were significantly correlated with higher occipital glutamate concentrations (*r*_24_ = −0.49, uncorrected *p* = 0.012; Bonferroni corrected *p* = 0.036; Figure 1G). Across the scalp, we observed a posterior group of 10 neighboring electrodes for which aperiodic slopes correlated significantly with occipital glutamate (cluster corrected *p* < 0.04; Figure 1H) and one additional electrode (T8) that did not survive cluster correction (cluster corrected *p* = 0.5; Figure 1H; gray dot).

Likewise, we examined the association between aperiodic slope (eyes open) and occipital GABA levels in the same participants. As GABA is thought to be a marker for cortical inhibition, we expected higher (more steep) aperiodic slopes would correlate with higher GABA concentrations, reflecting greater inhibitory tone. No significant correlation between GABA levels and aperiodic slope across occipital electrodes was found (*r*_24_ = 0.12, uncorrected *p* = 0.5; Supplemental Figure 2A), nor indeed for any of the electrode locations (uncorrected *p*-values > 0.05; Supplemental Figure 2B).

To test whether the relationship between aperiodic slope and metabolite concentration was specific to glutamate, we also examined correlations for NAA, a metabolite thought to reflect neuronal integrity ^53,54^, which is not thought to be closely linked to excitatory or inhibitory neural functions ^28,55^. The correlation between NAA levels and the aperiodic slope (eyes open) from occipital electrodes was not significant (*r*_24_ = −0.22, uncorrected *p* = 0.2; Supplemental Figure 3A). When examining the relationship between NAA and slope across the scalp, there were 4 groups of electrodes that did not pass cluster correction (cluster corrected *p*-values ≥ 0.12; Supplemental Figure 3B).As we scaled our metabolite measurements relative to water, we examined whether the observed relationship between glutamate and aperiodic slope might be attributable to individual differences in the water signal.

However, we saw no significant correlation between aperiodic slope across occipital electrodes and the area of the unsuppressed water peak (*r*_24_ = −0.01, uncorrected *p* = 0.9; Supplemental Figure 3C and D). Likewise, we saw no significant correlations between aperiodic slope and either the SNR (*r*_24_ = −0.05, uncorrected *p* = 0.8; Supplemental Figure 3E and F) or linewidth (*r*_24_ = −0.28, uncorrected *p* = 0.17; Supplemental Figure 3G and H) of our MRS data, suggesting individual differences in data quality were unlikely to explain the correlation with glutamate reported above.

Finally, we examined whether the observed correlation between glutamate and aperiodic slope might have been driven by individual differences in age, sex assigned at birth, time of day of the MRS scan, or VOI tissue composition (i.e., gray matter fraction). Here, we quantified the relationship between glutamate levels and aperiodic slopes using a linear mixed-effects model with the above factors included as covariates. The relationship between slope and glutamate concentration remained significant (linear mixed-effects model, parameter estimate [SE] = −0.15 [0.06], *t*_20_ = −2.6, *p* = 0.016) even when accounting for these factors. We did not find a significant association between aperiodic slope and any of the above covariates; age (parameter estimate [SE] = −0.005 [0.003], *t*_20_ = −1.7, *p* = 0.11), sex assigned at birth (parameter estimate [SE] = −0.048 [0.075], *t*_20_ = −0.65, *p* = 0.5), time of day of MRS scan (parameter estimate [SE] = −0.0002 [0.0002], *t*_20_ = −0.77, *p* = 0.4), and GM fraction (parameter estimate [SE] = 0.004 [0.82], *t*_20_ = 0.005, *p* = 0.9).

## Discussion

To the best of our knowledge, our study is the first to show that individual differences in resting aperiodic slope are related to glutamate concentration in the occipital cortex. Our results extend the findings of Gao et al. ^10^ and others ^56–58^ showing that the aperiodic slope of electrophysiological power spectra reflects the local balance of E:I. We show that for human EEG data, the aperiodic slope is related, in part, to excitatory tone, as indexed by glutamate concentrations from MRS.

Our analyses focused on occipital brain areas associated with visual processing. We used a visual stimulus as a functional localizer to place our MRS VOI within the occipital cortex, centered over the calcarine sulcus. Likewise, we used an EEG electrode montage with a higher density of electrodes over posterior scalp regions, and focused our analyses, in part, on averaged data from occipital electrodes. Both EEG and MRS data were collected in the absence of salient visual stimulation, suggesting that our results are more reflective of a baseline brain state rather than specific to visual processing. However, others have reported changes in aperiodic slope related to visual task performance ^11,59^ and have found that such changes may be specific to visual brain areas ^56^. Within our EEG data, there was a cluster of 10 electrodes across the posterior part of the scalp that were significantly correlated with glutamate concentrations measured within the occipital MRS VOI. Because we used an average electrode reference for data collected from channels skewed toward posterior scalp regions, and given that EEG signals are spatially diffuse, it is difficult to assess the extent to which the observed relationship between aperiodic slope and glutamate concentration is specific to occipital brain regions.

While we predicted that the concentration of GABA would correlate with aperiodic slope, with more steep slopes indicative of higher GABA concentrations, we found this correlation was not significant. GABA is present at fairly low concentrations (∼1 mM) in the human brain, at a roughly ten times lower concentration than glutamate, which raises the question of whether we may have had insufficient statistical power to detect a relationship with GABA. On the other hand, our MRS data were collected at 7 T, which allowed us to quantify GABA separately from macromolecules since we used a basis set that included experimentally measured macromolecules ^48^. In addition, we found our GABA measurements were highly reliable as shown with test-retest data (Supplemental Figure 4; also see Genovese et al. ^60^), suggesting our methods are sufficient to measure small but reliable differences in GABA levels across individuals. Nevertheless, given that we saw no significant correlation between aperiodic slope and GABA levels, we are unable to make any strong conclusions about this relationship.

In conclusion, the observed relationship between aperiodic slope as derived from EEG, and glutamate concentration as derived from MRS, suggests that these methods provide similar indexes of cortical excitatory tone at rest. Our results are valuable given the interest in aperiodic slope as a measure of E:I balance ^10,11,14,56–58,61,62^. Developing tools to investigate excitation and inhibition in the human brain provides insight into the neural basis of brain function and neuropsychiatric conditions. Compared to MRS, EEG is more widely available, accessible to a greater proportion of participants, lower cost, and more easily deployed outside the laboratory. The congruence across these two measures supports the use of aperiodic slope as a non-invasive marker for glutamatergic tone in the human brain.

## Acknowledgements

We thank Kyle W. Killebrew for support with experimental design and data collection. This work was supported by funding from the National Institutes of Health: K01 MH120278, P41 EB015894, P30 NS076408, and UL1 TR002494.

## References

1. Bridi, M. et al. Daily Oscillation of the Excitation-Inhibition Balance in Visual Cortical Circuits. Neuron 105, 621–629.e4 (2020).

2. Vyazovskiy, V. V., Cirelli, C., Pfister-Genskow, M., Faraguna, U. & Tononi, G. Molecular and electrophysiological evidence for net synaptic potentiation in wake and depression in sleep. Nat. Neurosci. 11, 200–208 (2008).

3. Adesnik, H. Layer-specific excitation/inhibition balances during neuronal synchronization in the visual cortex. J. Physiol. 596, 1639–1657 (2018).

4. Bateup, H. S. et al. Excitatory/inhibitory synaptic imbalance leads to hippocampal hyperexcitability in mouse models of Tuberous Sclerosis. Neuron 78, 510–522 (2013).

5. Wallace, M. L., Burette, A. C., Weinberg, R. J. & Philpot, B. D. Maternal loss of Ube3a produces an excitatory/inhibitory imbalance through neuron type-specific synaptic defects. Neuron 74, 793–800 (2012).

6. Monier, C., Fournier, J. & Frégnac, Y. In vitro and in vivo measures of evoked excitatory and inhibitory conductance dynamics in sensory cortices. J. Neurosci. Methods 169, 323–365 (2008).

7. Xue, M., Atallah, B. V. & Scanziani, M. Equalizing Excitation-Inhibition Ratios across Visual Cortical Neurons. Nature 511, 596–600 (2014).

8. Donoghue, T. et al. Parameterizing neural power spectra into periodic and aperiodic components. Nat. Neurosci. 23, 1655–1665 (2020).

9. He, B. J. Scale-free brain activity: past, present and future. Trends Cogn. Sci. 18, 480–487 (2014).

10. Gao, R., Peterson, E. J. & Voytek, B. Inferring synaptic excitation/inhibition balance from field potentials. NeuroImage 158, 70–78 (2017).

11. Deodato, M. & Melcher, D. Aperiodic EEG Predicts Variability of Visual Temporal Processing. J. Neurosci. 44, (2024).

12. KaŁamaŁa, P. et al. Event-induced modulation of aperiodic background EEG: Attention-dependent and age-related shifts in E:I balance, and their consequences for behavior. Imaging Neurosci. 2, 1–18 (2024).

13. Finley, A. J. et al. Resting EEG Periodic and Aperiodic Components Predict Cognitive Decline Over 10 Years. BioRxiv Prepr. Serv. Biol. 2023.07.17.549371 (2024) doi:10.1101/2023.07.17.549371.

14. McKeon, S. D. et al. Aperiodic EEG and 7T MRSI evidence for maturation of E/I balance supporting the development of working memory through adolescence. Dev. Cogn. Neurosci. 66, 101373 (2024).

15. Merkin, A. et al. Do age-related differences in aperiodic neural activity explain differences in resting EEG alpha? Neurobiol. Aging 121, 78–87 (2023).

16. Voytek, B. et al. Age-Related Changes in 1/f Neural Electrophysiological Noise. J. Neurosci. Off. J. Soc. Neurosci. 35, 13257–13265 (2015).

17. Pani, S. M., Saba, L. & Fraschini, M. Clinical applications of EEG power spectra aperiodic component analysis: A mini-review. Clin. Neurophysiol. Off. J. Int. Fed. Clin. Neurophysiol. 143, 1–13 (2022).

18. Peterson, E. J., Rosen, B. Q., Belger, A., Voytek, B. & Campbell, A. M. Aperiodic Neural Activity is a Better Predictor of Schizophrenia than Neural Oscillations. Clin. EEG Neurosci. 54, 434–445 (2023).

19. Lea-Carnall, C. A., El-Deredy, W., Stagg, C. J., Williams, S. R. & Trujillo-Barreto, N. J. A mean-field model of glutamate and GABA synaptic dynamics for functional MRS. NeuroImage 266, 119813 (2023).

20. Ip, I. B. & Bridge, H. Investigating the neurochemistry of the human visual system using magnetic resonance spectroscopy. Brain Struct. Funct. 227, 1491–1505 (2022).

21. Schallmo, M.-P. et al. Glutamatergic facilitation of neural responses in MT enhances motion perception in humans. NeuroImage 184, 925–931 (2019).

22. Conti, F. & Weinberg, R. J. Shaping excitation at glutamatergic synapses. Trends Neurosci. 22, 451–458 (1999).

23. de la Vega, A. et al. Individual Differences in the Balance of GABA to Glutamate in pFC Predict the Ability to Select among Competing Options. J. Cogn. Neurosci. 26, 2490–2502 (2014).

24. Kumar, J. et al. Glutathione and glutamate in schizophrenia: a 7T MRS study. Mol. Psychiatry 25, 873–882 (2020).

25. Willis, H. E. et al. GABA and Glutamate in hMT+ Link to Individual Differences in Residual Visual Function After Occipital Stroke. Stroke 54, 2286–2295 (2023).

26. Apšvalka, D., Gadie, A., Clemence, M. & Mullins, P. G. Event-related dynamics of glutamate and BOLD effects measured using functional magnetic resonance spectroscopy (fMRS) at 3T in a repetition suppression paradigm. NeuroImage 118, 292–300 (2015).

27. Bednařík, P. et al. Neurochemical and BOLD responses during neuronal activation measured in the human visual cortex at 7 Tesla. J. Cereb. Blood Flow Metab. Off. J. Int. Soc. Cereb. Blood Flow Metab. 35, 601–610 (2015).

28. Chen, C. et al. Activation induced changes in GABA: Functional MRS at 7T with MEGA-sLASER. NeuroImage 156, 207–213 (2017).

29. Ip, I. B. et al. Combined fMRI-MRS acquires simultaneous glutamate and BOLD-fMRI signals in the human brain. NeuroImage 155, 113–119 (2017).

30. Kurcyus, K. et al. Opposite Dynamics of GABA and Glutamate Levels in the Occipital Cortex during Visual Processing. J. Neurosci. Off. J. Soc. Neurosci. 38, 9967–9976 (2018).

31. Mangia, S. et al. Sustained neuronal activation raises oxidative metabolism to a new steady-state level: evidence from 1H NMR spectroscopy in the human visual cortex. J. Cereb. Blood Flow Metab. Off. J. Int. Soc. Cereb. Blood Flow Metab. 27, 1055–1063 (2007).

32. Mullins, P. G., Rowland, L. M., Jung, R. E. & Sibbitt, W. L. A novel technique to study the brain’s response to pain: proton magnetic resonance spectroscopy. NeuroImage 26, 642–646 (2005).

33. Koolschijn, R. S. et al. Memory recall involves a transient break in excitatory-inhibitory balance. eLife 10, e70071 (2021).

34. Stanley, J. A. et al. Functional dynamics of hippocampal glutamate during associative learning assessed with in vivo 1H functional magnetic resonance spectroscopy. NeuroImage 153, 189–197 (2017).

35. Mullins, P. G. Towards a theory of functional magnetic resonance spectroscopy (FMRS): A meta-analysis and discussion of using MRS to measure changes in neurotransmitters in real time. Scand. J. Psychol. 59, 91–103 (2018).

36. Koolschijn, R. S., Clarke, W. T., Ip, I. B., Emir, U. E. & Barron, H. C. Event-related functional magnetic resonance spectroscopy. NeuroImage 276, 120194 (2023).

37. Chatrian, G. E., Lettich, E. & and Nelson, P. L. Ten Percent Electrode System for Topographic Studies of Spontaneous and Evoked EEG Activities. Am. J. EEG Technol. 25, 83–92 (1985).

38. Chatrian, G. E., Lettich, E. & Nelson, P. L. Modified nomenclature for the ‘10%’ electrode system. J. Clin. Neurophysiol. Off. Publ. Am. Electroencephalogr. Soc. 5, 183–186 (1988).

39. Oostenveld, R. & Praamstra, P. The five percent electrode system for high-resolution EEG and ERP measurements. Clin. Neurophysiol. Off. J. Int. Fed. Clin. Neurophysiol. 112, 713–719 (2001).

40. Lin, A. et al. Minimum Reporting Standards for in vivo Magnetic Resonance Spectroscopy (MRSinMRS): Experts’ consensus recommendations. NMR Biomed. 34, e4484 (2021).

41. Tkác, I. et al. In vivo 1H NMR spectroscopy of the human brain at 7 T. Magn. Reson. Med. 46, 451–456 (2001).

42. Gruetter, R. Automatic, localized in vivo adjustment of all first- and second-order shim coils. Magn. Reson. Med. 29, 804–811 (1993).

43. Gruetter, R. & Tkác, I. Field mapping without reference scan using asymmetric echo-planar techniques. Magn. Reson. Med. 43, 319–323 (2000).

44. Delorme, A. & Makeig, S. EEGLAB: an open source toolbox for analysis of single-trial EEG dynamics including independent component analysis. J. Neurosci. Methods 134, 9–21 (2004).

45. Welch, P. The use of fast Fourier transform for the estimation of power spectra: A method based on time averaging over short, modified periodograms. IEEE Trans. Audio Electroacoustics 15, 70–73 (1967).

46. Provencher, S. W. Automatic quantitation of localized in vivo 1H spectra with LCModel. NMR Biomed. 14, 260–264 (2001).

47. Behar, K. L., Rothman, D. L., Spencer, D. D. & Petroff, O. A. Analysis of macromolecule resonances in 1H NMR spectra of human brain. Magn. Reson. Med. 32, 294–302 (1994).

48. Marjańska, M. et al. Region-specific aging of the human brain as evidenced by neurochemical profiles measured noninvasively in the posterior cingulate cortex and the occipital lobe using 1H magnetic resonance spectroscopy at 7 T. Neuroscience 354, 168–177 (2017).

49. Giapitzakis, I.-A., Borbath, T., Murali-Manohar, S., Avdievich, N. & Henning, A. Investigation of the influence of macromolecules and spline baseline in the fitting model of human brain spectra at 9.4T. Magn. Reson. Med. 81, 746–758 (2019).

50. Marjańska, M., Deelchand, D. K., Kreis, R., & 2016 ISMRM MRS Study Group Fitting Challenge Team. Results and interpretation of a fitting challenge for MR spectroscopy set up by the MRS study group of ISMRM. Magn. Reson. Med. 87, 11–32 (2022).

51. Marjańska, M. & Terpstra, M. Influence of fitting approaches in LCModel on MRS quantification focusing on age-specific macromolecules and the spline baseline. NMR Biomed. e4197 (2019) doi:10.1002/nbm.4197.

52. Fischl, B. FreeSurfer. NeuroImage 62, 774–781 (2012).

53. Lin, A., Ross, B. D., Harris, K. & Wong, W. Efficacy of proton magnetic resonance spectroscopy in neurological diagnosis and neurotherapeutic decision making. NeuroRx J. Am. Soc. Exp. Neurother. 2, 197–214 (2005).

54. Rae, C. D. A guide to the metabolic pathways and function of metabolites observed in human brain 1H magnetic resonance spectra. Neurochem. Res. 39, 1–36 (2014).

55. DiNuzzo, M. et al. Perception is associated with the brain’s metabolic response to sensory stimulation. eLife 11, e71016 (2022).

56. Podvalny, E. et al. A unifying principle underlying the extracellular field potential spectral responses in the human cortex. J. Neurophysiol. 114, 505–519 (2015).

57. van Bueren, N. E. R., van der Ven, S. H. G., Hochman, S., Sella, F. & Cohen Kadosh, R. Human neuronal excitation/inhibition balance explains and predicts neurostimulation induced learning benefits. PLOS Biol. 21, e3002193 (2023).

58. Waschke, L. et al. Modality-specific tracking of attention and sensory statistics in the human electrophysiological spectral exponent. eLife 10, e70068 (2021).

59. Ouyang, G., Hildebrandt, A., Schmitz, F. & Herrmann, C. S. Decomposing alpha and 1/f brain activities reveals their differential associations with cognitive processing speed. NeuroImage 205, 116304 (2020).

60. Genovese, G., Deelchand, D. K., Terpstra, M. & Marjańska, M. Quantification of GABA concentration measured noninvasively in the human posterior cingulate cortex with 7 T ultra-short-TE MR spectroscopy. Magn. Reson. Med. 89, 886–897 (2023).

61. Gao, Y. et al. Catecholaminergic Modulation of Metacontrol Is Reflected in Aperiodic EEG Activity and Predicted by Baseline GABA+ and Glx Concentrations. Hum. Brain Mapp. 46, e70173 (2025).

62. Wiest, C. et al. The aperiodic exponent of subthalamic field potentials reflects excitation/inhibition balance in Parkinsonism. eLife 12, e82467 (2023).

