## Supplemental Information for "Aperiodic slope reflects glutamatergic tone in the human brain"

### Supplemental Methods

*Participants*

Participants were between 18 - 60 years old (EEG n = 35, mean age 39 years ± 11 SD, 19 males, 16 females; MRS n = 27, mean age 39 years ± 11 SD, 14 males, 12 females). Gender identity was collected and did not differ from sex assigned at birth for this cohort. Participants spoke English as their primary language, did not have a learning disability or an IQ lower than 70, had no current substance dependence (besides nicotine), current or past central nervous system disease, history of traumatic brain injury (history of skull fracture or loss of consciousness longer than 30 minutes), electroconvulsive therapy within the last year, tardive dyskinesia, visual or hearing impairment, or history of psychotic or affective disorders as assessed with the Mini International Neuropsychiatric Interview for Psychotic Disorders Studies (version 7.0.2). Participants were assessed with a Snellen visual acuity chart on the day of data collection (both EEG and MRS visits) and confirmed to have normal (or corrected-to-normal) binocular vision (visual acuity of 20/40; decimal fraction = 0.5, or better). Participants were able to fit into the MRI scanner comfortably.

Demographics

| | EEG | EEG + MRS |
|---|---|---|
| **N** | 35 | 26 |
| **Age (years)** | 39 ± 11 | 38 ± 11 |
| **Sex assigned at birth** | 16 F; 19 M | 12 F; 14 M |
| **Race** | 5 Asian; 2 Black; 25 White; 3 More than one race | 2 Asian; 1 Black; 23 White |
| **Ethnicity** | 2 Hispanic; 33 Not Hispanic | 26 Not Hispanic |
| **Education (years)** | 16 ± 2 | 16 ± 2 |

Supplemental Table 1. Demographic information for participants included with EEG results (first column) and a subset of participants with EEG and MRS (second column).

*Apparatus*

EEG

The EEG montage contained the following electrodes: (Fp1, Fp2, F7, Fz, F8, FC5, FC6, T7, T8, Cz, CP5, CPz, CP6, TP9, TP10, P7, P3, Pz, P4, P8, PO7, PO3, POz, PO4, PO8, O1, Oz, O2, PO9, PO10, O9, O10)

During setup, the scalp was abraded to reduce electrode impedances below 10 kΩ. Data were referenced online to Cz and collected at a sampling rate of 1000 Hz using a 24-bit actiCHamp amplifier (no online filter) and BrainVision Recorder software (Version 1.22.0002; Copyright 2000-2019 Brain Products GmbH). Head position was maintained using a fixed chinrest (60 cm viewing distance from screen) and participants were viewed from an adjacent experimenter room using an infrared camera to monitor for motion and compliance with task instructions.

MRS

The 7 T scanner (software version VE12U) was equipped with an 8-kW RF power amplifier and body gradients with 70 mT/m maximum amplitude and 200 T/m/s maximum slew rate. The 3 T scanner was equipped with gradients with 200 T/m/s maximum slew rate and 80 mT/m maximum amplitude.

Participants were instructed to minimize movement and were padded with head, neck, and lumbar pillows to increase their comfort throughout the scan. Head motion was monitored during scanning with an in-scanner motion tracking system from Metria Innovation Inc. (Milwaukee, WI), which used a Moiré Phase Tracking marker that was attached to the participant's nose. If motion exceeded > 5 mm from starting position, the MRS scan was discontinued and repeated.

*Average VOI placement*

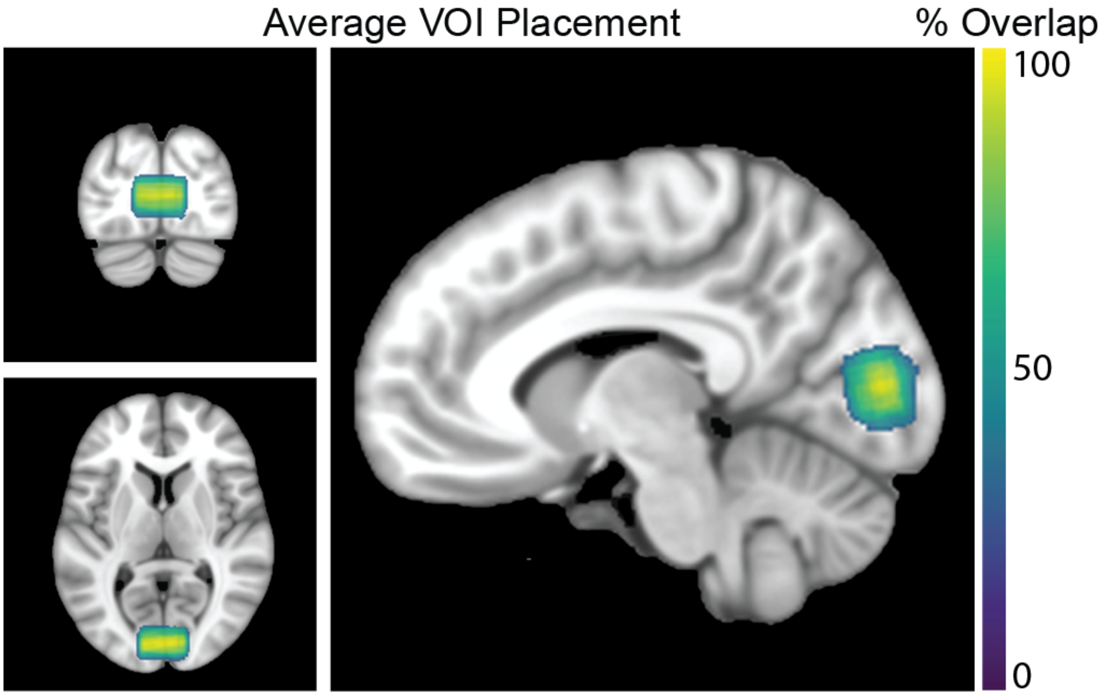


Supplemental Figure 1. The average occipital VOI placement for all the participants included in the MRS & EGG analysis (n = 26). Color bar shows percentage overlap of VOI.

*Experimental protocol*

*EEG*

The EEG resting task was presented on an Asus VG248QE monitor using a PC (Windows 10). During eyes open blocks, participants were instructed to fixate on a small (0.4°) blue central fixation circle on a black screen (luminance 0.3 cd/m$^2$).

*MRS*

STEAM sequence parameters were as follows: echo time (TE) = 8 ms, repetition time (TR) = 5 s, mixing time (TM) = 32 ms, chemical shift displacement error = 4%, spectral bandwidth = 6000 Hz, number of complex data points = 2048, transmitter frequency = 3 ppm, hsinc pulse = 1.28 ms. Gradient echo fMRI data were obtained for the localizer using the following parameters: TR = 3000 ms, TE = 22.2 ms, echo spacing = 0.73 ms, flip angle = 45°, resolution = 1.6 mm$^3$, partial Fourier = 5/8, 30 coronal slices, field of view = 200 × 200 mm$^2$, phase encode direction = right-left.

Water suppression was performed by acquiring 4 free induction decays (FIDs; averages) and checking that the water peak was lower than the *N*-acetyl-aspartate (NAA) peak. In cases where water suppression failed, we adjusted the VAPOR parameters manually. The transmit voltage was adjusted for each VOI to calibrate the $B_1$ field for both the 90° excitation pulse and the water suppression flip angles.

Anatomical images were collected in a separate session at 3 T with the following parameters: $T_1$: TE = 1.81, 3.6, 5.39 and 7.18 ms, TR = 2500 ms, flip angle = 8°, resolution = 0.8 $mm^3$, echo spacing = 11.2 ms, 208 sagittal slices, FOV = 240 x 256 $mm^2$, partial Fourier = 6/8. $T_2$: TE = 564 ms, TR = 3200 ms, variable flip angle, resolution = 0.8 $mm^3$, echo spacing = 3.86 ms, 208 sagittal slices, FOV = 240 x 256 $mm^2$, no partial Fourier. The 7 T $T_1$ images were acquired with the following parameters: TE = 3.2 ms, TR = 2500 ms, flip angle = 5°, resolution = 1.3 $mm^3$, echo spacing = 7.9 ms, 64 sagittal slices, FOV = 160 x 160 $mm^2$, partial Fourier = 6/8.

_MRS task_

Stimuli were presented in PsychoPy (version 1.85.2; [1] on an acrylic screen; 3M Vikuiti, Maplewood, MN) at the back of the scanner bore, using an NEC projector (60 Hz refresh rate; mean luminance = 290 cd/$m^2$). The screen was viewed from a distance of 100 cm via a mirror mounted to the head coil. Participants were instructed to maintain fixation on a small (0.2°) central square presented on a mean gray background and, on each trial, judge which of two small dots (0.125° radii, presented 0.3° directly to left and right of fixation) was brighter. Stimulus contrast ranged from 0.8 - 25% on a 3-down-1-up staircase, with a starting contrast of 4.7% using a task designed to converge at a contrast level for which each participant's discrimination accuracy was 79% [2]. Data were collected in 5-minute blocks, which alternated with 5 minutes of the same task, but with the addition of a large contrast reversing grating stimulus (data from this portion of the task will be presented elsewhere). The stimulus consisted of a ring-shaped contrast reversing grating stimulus presented at the center of the screen with the following parameters: inner radius = 0.75°, outer radius = 8°, contrast = 80%, spatial frequency = 1.1 cycles / °, temporal frequency = 4 Hz, stimulus duration = 0.75 s, inter-stimulus interval = 0.75 s. Grating orientation was randomized across trials. The data presented here were acquired without the large visual stimulus (i.e., fixation task and gray background only). For the localizer, stimuli were presented in 12 s blocks, which alternated with 12 blank blocks (i.e., fixation task only, no gratings). There were 8 stimulus blocks and 9 blank blocks in the functional localizer, with a scan duration of 3.4 min.

*Data processing and analysis*

_EEG_

EEG filters used the following parameters: high-pass filtered at 0.05 Hz (6 dB/octave roll-off) and notch filtered (58 - 62 Hz; 243 db/octave roll-off) to remove AC electrical noise, using a Hamming windowed sinc finite impulse response filter. Independent component analysis was performed using the infomax algorithm [3].

_MRS_

After frequency and phase correction, we used an automated method to identify FIDs for which artifacts remained. Specifically for each FID, we calculated the Pearson correlation across frequencies between the spectral data for that FID and the median of all other FIDs. FIDs for which the correlation values were substantially lower than average (i.e., greater than 10 median absolute deviations from the median) were excluded. Frequency and phase correction were then repeated.

The basis set included 18 metabolites: ascorbic acid, aspartic acid, creatine, GABA, glucose, glutamate, glutamine, glutathione, glycerophosphorylcholine, lactate, *myo*-inositol, NAA, *N*-acetyl-aspartylglutamate, phosphocreatine, phosphorylcholine, phosphorylethanolamine, *scyllo*-inositol, taurine, lipids and macromolecules.

The 7 T $T_1$-weighted anatomical images were aligned to the whole brain 3 T $T_1$-weighted anatomical images, with an average registration for all participants shown in Supplemental Figure 1. The tissue segmentations were used to quantify the tissue fraction within each VOI for each participant (example participant in Figure 1E in the main text), and metabolite concentrations were corrected for individual tissue fraction in MATLAB (2014a). The metabolite concentrations were not corrected for $T_1$ and $T_2$ relaxation times, as these effects are thought to be nominal with long TRs (5 s) and ultra-short TEs (8 ms).

SNR is quantified in LCModel using the maximum metabolite signal value (i.e., the NAA peak) divided by two times the mean square root of the residuals. After data quality checks, we excluded 5 out of 92 runs across all participants, and we removed 1 participant who had less than 2 runs after exclusions.

<u>*Water scaling*</u>

| | **Water content** | **$T_1$ relaxation time constant (ms)** | **$T_2$ relaxation time constant (ms)** |
|---|---:|---:|---:|
| **GM** | 80% | 2130 | 50 |
| **WM** | 71% | 1220 | 55 |
| **CSF** | 97% | 4425 | 141 |

Supplemental Table 2. Correction parameters for water scaling. Columns represent the water content (%), $T_1$ and $T_2$ relaxation time constants (ms) for the cortical tissues of gray matter (GM), white matter (WM) and cerebrospinal fluid (CSF; rows). These are based on values from Marjańska *et al.* [4] .

*Statistical analysis*

We corrected for multiple comparisons by performing permutation tests using Monte Carlo simulations (n = 10,000 iterations per metabolite) in which data were permuted randomly across participants. For each iteration, we used the same random assignment for data across all electrodes, to preserve spatial autocorrelations in the EEG data [5,6] . Neighboring electrodes were decided by consensus among three authors (A.S., H.M., M-P.S.). For each iteration, we performed paired *t*-tests (for eyes open vs. eyes closed, 1-tailed) or Spearman correlations (for associations with MRS data) for each electrode using the permuted data. This was done to define a null distribution for the number and size of clusters of significant neighboring electrodes at an uncorrected $p < 0.05$. The cluster corrected $p$ value was defined as the number of significant clusters from the permuted data that were as large or larger than the clusters in the observed data.

A linear mixed-effects model was applied to assess the glutamate-slope relationship while accounting for age, sex assigned at birth, and normalized gray matter (GM) tissue fraction (Figure 1E in the main text). Tissue fraction was calculated by taking the percentage of GM divided by the sum of the percentage of WM and GM, according to the following equation:

$$GM\ fraction = \frac{\%\ GM}{(\%\ GM + \%\ WM)}$$

**Supplemental Results**

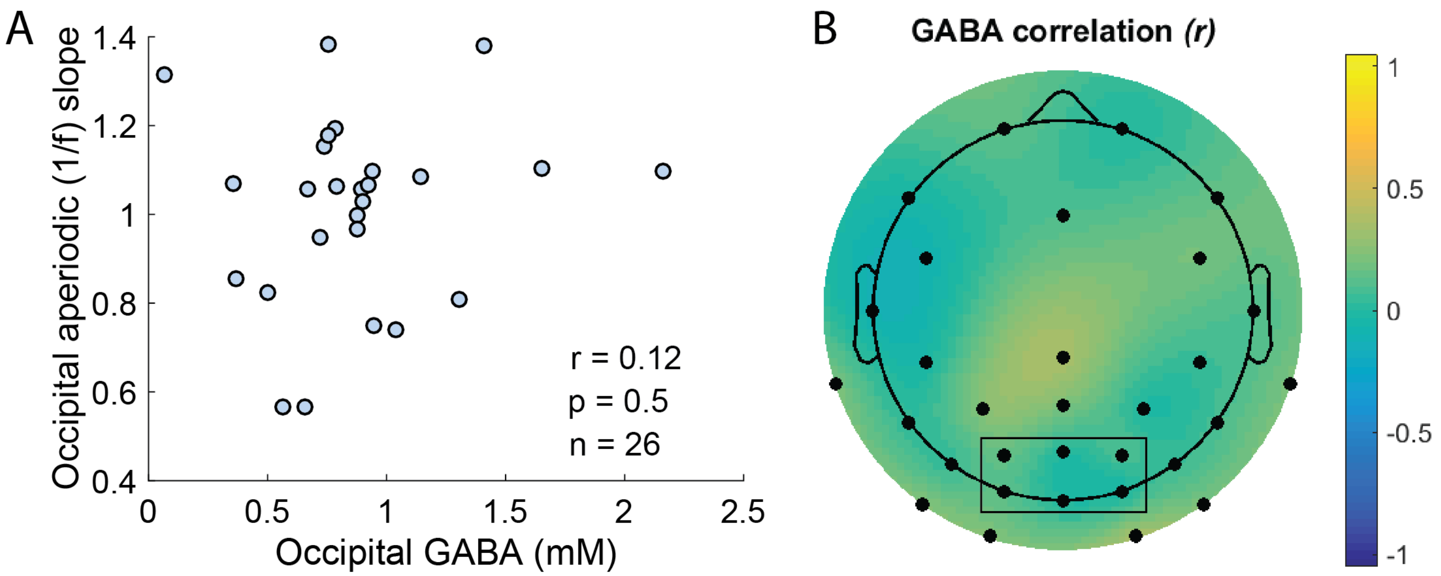


Supplemental Figure 2. A) Correlation between the average eyes open aperiodic 1/f slope from the 6 occipital electrodes (O1, Oz, O2, PO3, POz, PO4; black box) and GABA concentration (circles) is plotted. B) Topoplot showing the correlation between GABA concentration and eyes open aperiodic slope measured at each electrode. The interpolated color plot over the scalp shows the *r* value. Points show the electrode locations, red indicates a significant correlation (cluster corrected *p* < 0.05; none were significant).

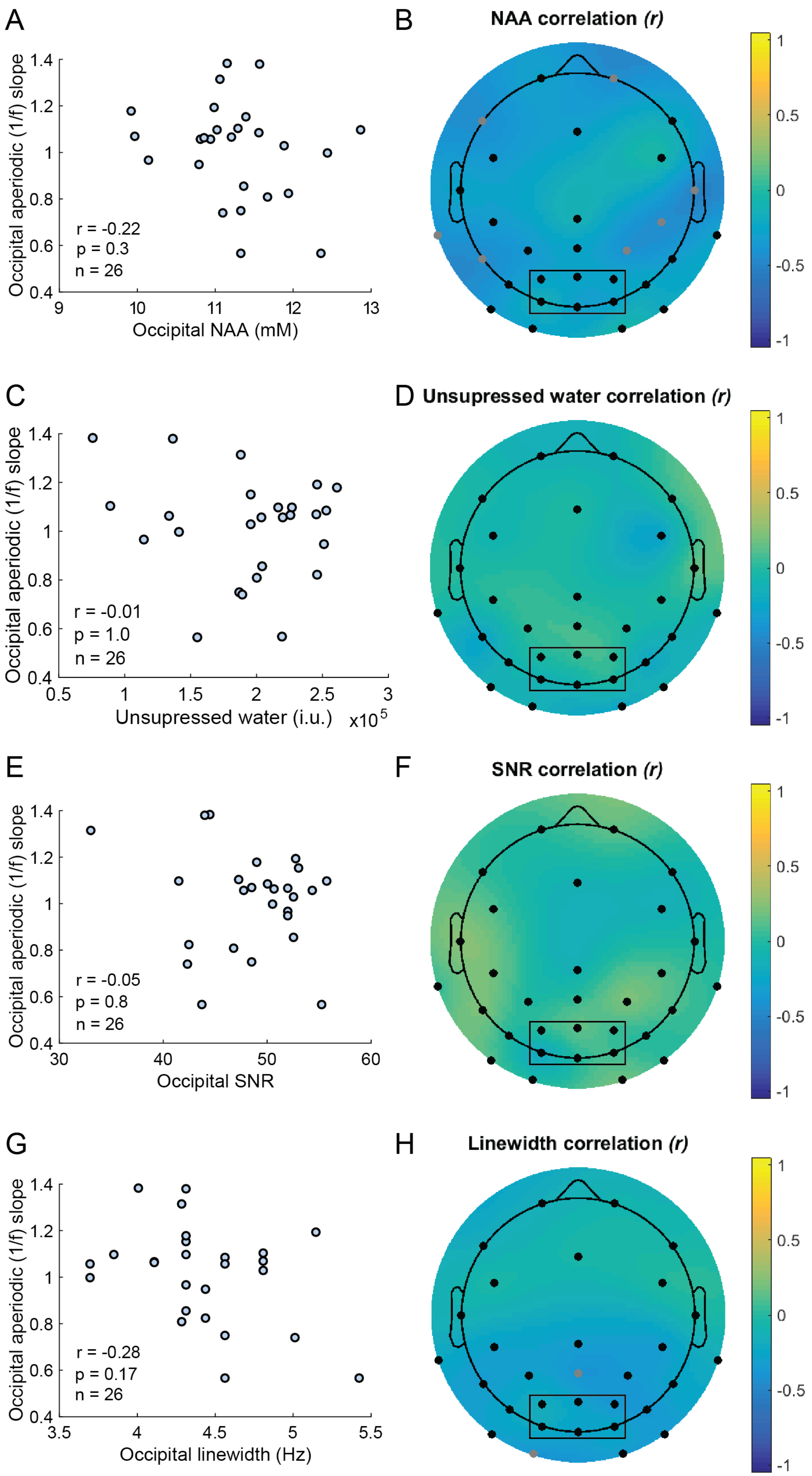
A
Occipital aperiodic (1/f) slope
r = -0.22
p = 0.3
n = 26
Occipital NAA (mM)
B
NAA correlation (r)
C
r = -0.01
p = 1.0
n = 26
Unsupressed water (i.u.)
x10^5
D
Unsupressed water correlation (r)
E
r = -0.05
p = 0.8
n = 26
Occipital SNR
F
SNR correlation (r)
G
r = -0.28
p = 0.17
n = 26
Occipital linewidth (Hz)
H
Linewidth correlation (r)

Supplemental Figure 3. A) Correlation between eyes open aperiodic 1/f slope from 6 occipital electrodes (O1, Oz, O2, PO3, POz, PO4; black box) and occipital NAA concentration. B) Topoplot showing the correlation between NAA concentration and eyes open aperiodic slope measured at each electrode. The interpolated color plot over the scalp shows the *r* value. Points show the electrode locations, red indicates a significant correlation (cluster corrected $p < 0.05$; none were significant), gray indicates electrodes that did not pass cluster correction (uncorrected $p < 0.05$). Repeated for the area of the unsuppressed water peak (C, D), SNR (E, F) and linewidth (G, H).

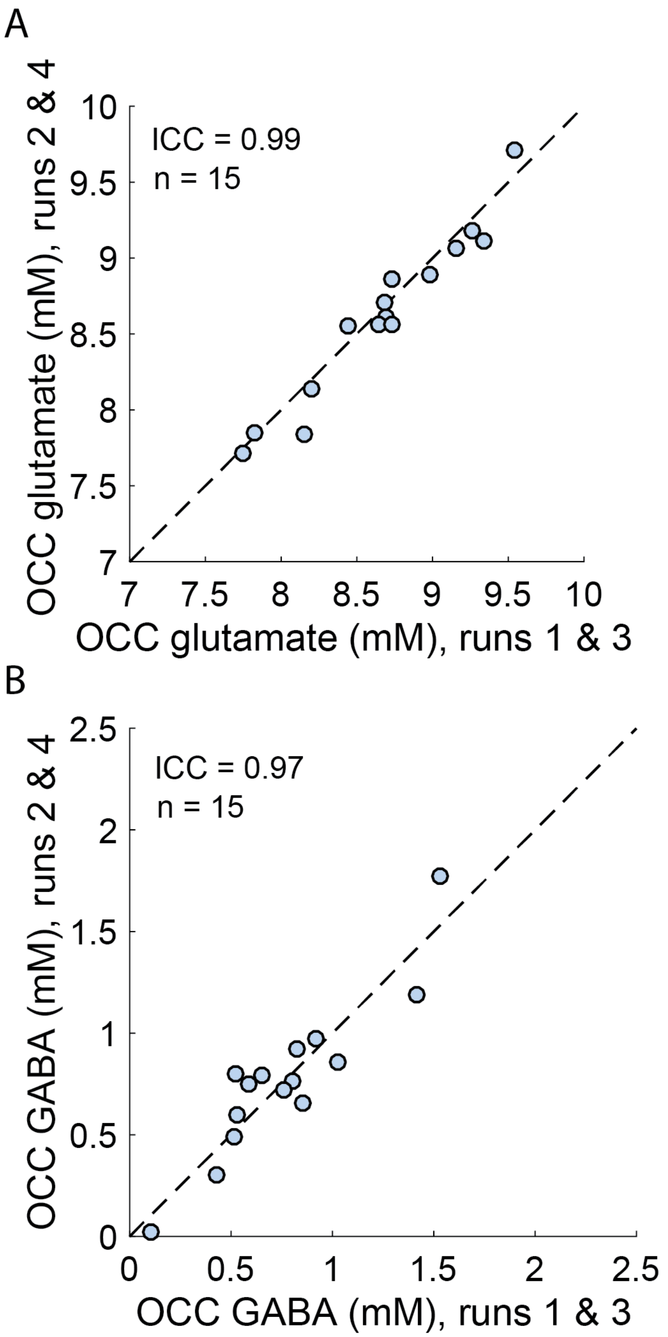


Supplemental Figure 4. Glutamate (A) and GABA (B) test-retest data from a subset of participants (n = 15) with 4 MRS scan runs. The average concentration from runs 1 & 3 and runs 2 & 4 are plotted, along with the unity line. Intra-class correlation coefficient was very high (Glutamate ICC = 0.99; GABA ICC: 0.97), indicating reliability.

### Supplemental References


1. Peirce, J. W. PsychoPy—Psychophysics software in Python. *J. Neurosci. Methods* **162**, 8–13 (2007).
2. García-Pérez, M. A. Forced-choice staircases with fixed step sizes: asymptotic and small-sample properties. *Vision Res.* **38**, 1861–1881 (1998).
3. Lee, T. W., Girolami, M. & Sejnowski, T. J. Independent component analysis using an extended infomax algorithm for mixed subgaussian and supergaussian sources. *Neural Comput.* **11**, 417–441 (1999).
4. Marjańska, M. *et al.* Region-specific aging of the human brain as evidenced by neurochemical profiles measured noninvasively in the posterior cingulate cortex and the occipital lobe using 1H magnetic resonance spectroscopy at 7 T. *Neuroscience* **354**, 168–177 (2017).
5. Bae, G.-Y. & Luck, S. J. Appropriate Correction for Multiple Comparisons in Decoding of ERP Data: A Re-Analysis of Bae & Luck (2018). 672741 Preprint at https://doi.org/10.1101/672741 (2019).
6. Eklund, A., Nichols, T. E. & Knutsson, H. Cluster failure: Why fMRI inferences for spatial extent have inflated false-positive rates. *Proc. Natl. Acad. Sci.* **113**, 7900–7905 (2016).
